# Preparation and characterization of phospholipid stabilized nanoemulsions in small-scale

**DOI:** 10.1101/603803

**Authors:** Shila Gurung, Martin Holzer, Sabine Barnert, Rolf Schubert

**Author notes:** Corresponding author; (SG).

## Abstract

Phospholipids have been used to prepare liposomes. The use of phospholipids to stabilize nanoemulsions may cause spontaneous formation of liposomes. The main objective of this study is to develop a method to prepare phospholipid stabilized nanoemulsions in small scale (< 1 mL) and to minimize the formation of liposomes.

A combination of hand extrusion and detergent removal methods was used in this study. Extrusion through polycarbonate membranes was performed in two steps, firstly using membranes of 400 nm followed by 200 nm membranes as the second step. Sodium cholate was used as a detergent to solubilize the formed liposomes which was later removed via dialysis. Nanoemulsions were characterized by measuring their particle size, polydispersity index and zeta-potential using Photon Correlation Spectroscopy and Cryo-TEM pictures. The stability of nanoemulsion stored under refrigeration was also studied.

Fifty-one extrusion cycles through polycarbonate membrane of 400 nm pore size followed by one-hundred fifty-three cycles through polycarbonate membrane of 200 nm produced nanoemulsions having particle size below 200 nm (diameter). The nanoemulsions were found to be homogenous as depicted by polydispersity index (PDI) value below 0.1. Similarly, the zeta-potential was measured to be above −30 mV which is sufficient to keep nanoemulsions stable for as long as 7 months when stored under refrigeration. The Cryo-TEM pictures revealed 30 mM to be an optimum concentration of sodium cholate to prepare homogenous nanoemulsions with negligible proportion of liposomes.

It was concluded that this method could be established as a small scale method of preparing nanoemulsions which will not only reduce the cost of preparation but also the disposal cost of toxic chemicals used for functionalizing nanoemulsions for scientific research.

## Introduction

Nanoemulsions are transparent or translucent systems generally covering a size range between 20-500 nm. Due to the small droplet size, the Brownian motion is sufficiently high to overcome the phenomena of physical destabilization caused by gravitational separation, flocculation and/or coalescence [1-4]. When the maximum droplet size of an emulsion is below 80 nm, it gains advanced properties compared to conventional emulsions, such as optical transparency, high colloidal stability and large interfacial area to volume ratio [5]. Nanoemulsions differ from microemulsions with respect to stability, preparation methods, dilution behavior and temperature fluctuations. In addition, nanoemulsions are thermodynamically unstable, but kinetically stable systems and require less surfactant than microemulsions. They are sometimes referred to as ‘approaching thermodynamic stability [5, 6].

Nanoemulsions have been extensively investigated as a promising drug delivery system for poorly water soluble substances. They have been used in intravenous, oral and ocular drug administrations to reduce drug side effects and improve the pharmacological effects of the loaded drugs [3].

Parenteral emulsions provide a number of potential advantages as drug delivery vehicle, such as reduction in pain, irritation and thrombophlebitis, reduced toxicity, improved stability and solubility, and the option for a targeted drug delivery approach [7]. The droplets remain stable under the conditions of temperature changes and/or dilutions and do not differ markedly from bulk values. This is especially advantageous for nanoemulsions because when they are injected into the bloodstream, changes in temperature, pH values, osmolarity etc. are likely to occur, affecting the physical properties of the loaded drug [8, 9].

Emulsion formation is a non-spontaneous process and therefore requires high energy [5, 10]. The basic composition of an emulsion system is oil, water and emulsifier. For the oil phase, long chain triglycerides such as triolein, soybean oil, safflower oil, sesame oil, and castor oil are approved for clinical use. Medium chain triglycerides are used alone or in combination and the approved ones include fractionated coconut oil, Miglyol^®^ 810 and 812, Neobee^®^, and Captex^®^ 300 [7]. Generally, the oil phase does not exceed 30% (w/w), due to which the application of emulsions in drug delivery is limited [11].

Emulsifier plays major role in the formation of nanoemulsions by lowering the interfacial tension and hence less stress is needed to break up a droplet [5, 12]. Depending upon the nature of the emulsifier used, they form an interfacial film at the o/w interface, which provides a mechanical barrier and repulsive force to stabilize the emulsion system. The repulsive forces provided by the emulsifiers can be electrostatic (e.g. lecithin), steric (e.g. block copolymer like Poloxamer 188) or electrosteric (a combination of both lecithin and block copolymers). Unfortunately, only a limited number of emulsifiers are approved and recognized as safe by the regulatory authorities [7, 13].

Most commonly used methods for preparing emulsions are simple pipe flow, static mixers and general stirrers, high speed mixers, colloid mills, high pressure homogenizers and ultrasonication. Other methods include the low energy emulsification method at constant temperature [4, 14] and the phase inversion temperature (PIT) [5, 15]. However, for preparing nanoemulsions, one is limited to the higher energy sources like high pressure homogenizer and ultrasonication [5, 16]. These so-called high energy methods supply enough energy to increase the interfacial area for generating submicronic droplets [8, 17].

Depending upon the preparation method, the size of nanoemulsions varies. Those prepared using PIT are relatively polydisperse and generally give higher Ostwald ripening rates than those prepared by high pressure homogenisation techniques [5]. By using ultrasonic devices having frequencies from 20 kHz to 1.0 MHz droplets typically between 100 and 1000 nm in diameter can be prepared. Therefore, they are mostly milky in appearance. However, use of a higher frequency of 2.4 MHz, could produce clear and transparent emulsions which are stable for 12 months even under surfactant-free conditions [18]. The main disadvantages of these high energy methods are difficulties for small scale preparation and therefore expensive.

In this study, we investigated a method to prepare stable and homogenous nanoemulsions having definite size and narrow size distribution (PDI below 0.1) in a small scale. This method would be very useful and cost effective in scientific research for the following reasons: a) for in vitro studies, very less volume of nanoemulsion (< 1 ml) is sufficient, b) to minimize the sample waste and disposal cost, c) to simplify the working set up by avoiding the use of compressed air (like in high pressure homogenizers), and d) to minimize the cost of preparation, especially when expensive chemicals like isotopes, fluorescence markers, nanoparticles surface modifiers like anchors and ligands or antibodies are used.

## Materials and Methods

### Materials

Medium chain triglyceride (Miglyol^®^ 812) was obtained from Sasol GmbH, Witten, Germany. Egg phospholipid (Lipoid^®^ E80) and Poloxamer 188 (Lutrol^®^ F68) were generously gifted by Lipoid GmbH, Ludwigshafen, Germany and BASF, Ludwigshafen, Germany, respectively. Glycerol (purity >99 %) and sodium cholate hydrate were purchased from Sigma-Aldrich Life Science, Steinheim, Germany. Similarly cholic acid (2, 4-^3^H) was obtained from American Radiolabeled Chemicals, Inc. St. Louis, MO, USA and phosphatidylcholine (DPPC, L-a-dipalmitoyl [1-palmitoyl-1-^14^C] from PerkinElmer, Inc. Albany Street, Boston, USA. Deionized water (18.2 MΩ.cm) was used for all dilutions. Slide-A-Lyzer^®^ (10 kDa membrane cut off) was commercially available from Thermo Scientific, Rockford, USA. All other chemicals were of analytical grade.

### Preparation of nanoemulsions

#### Hand extruded nanoemulsions

Crude emulsion was prepared by phase inversion temperature method. Briefly, the ingredients of aqueous phase (Table 1) were mixed in a round bottomed flask at 70 °C for about 30 min and 700 rpm using magnetic stirrer (MR 3001, Heidolph Instruments, Schwabach, Germany). Similarly, the oily phase was also prepared by mixing the ingredients (Table 1) in similar conditions. The aqueous phase was then added slowly to the oil phase upon constant stirring (700 rpm) but at an increased temperature of 80 °C. The mixture appeared apparently transparent in the beginning and then turned into translucent upon further addition of remaining aqueous phase. This translucent preparation was cooled for about 1 h in ice bath and was termed as crude emulsion. The theoretical concentration of phospholipid was maintained at 15.75 mM (1.2 %, w/w) throughout the study.

**Table 1.** Composition of nanoemulsion.

| Oil Phase | Aqueous Phase |
| --- | --- |
| Miglyol® (10 %, w/w) | Poloxamer 188 (1.0 %, w/w) |
| Phospholipid E80® (1.2 %, w/w) | Glycerol (2.5 %, w/w) |
|  | Deionized water (~ 86 %, w/w) |
|  | Sodium cholate (20 – 50 mM) |

Then the crude emulsion was extruded through polycarbonate membranes pre-equilibrated in deionized water for 15 – 30 min using a LiposoFast Basic device (Avestin Inc., Ottawa, Canada). The extrusion was done in two steps in a water bath at 65 °C. In the first step, the crude emulsion was extruded for 51 cycles (400 nm pore size) which was followed by additional 153 extrusion cycles (200 nm pore size). The membrane was changed after every 51 cycles to ensure that the membrane is not damaged. The odd number of extrusion cycle was used to collect the nanoemulsion from the second syringe so that the unextruded larger particles remained in the first syringe.

#### Nanoemulsions with sodium cholate

It is likely that nanoemulsions stabilized with phospholipids may contain liposomes. Therefore, detergents are used to solubilize lipid membranes [19] because at sufficient concentration, detergents such as sodium cholate solubilize the phospholipid to form mixed micelles (MMs). When the amount of detergent in the MMs is reduced further, the MMs enlarge to form liposomes [20, 21]. Thus, different concentrations (20 – 50 mM) of sodium cholate were used to solubilize the liposomes formed during the preparation of nanoemulsions. The concentration of sodium cholate influences the formation and solubilization of liposomes. The sodium cholate was added to the aqueous phase and the crude emulsion was prepared as described previously.

The detergent was removed by dialysis as in the preparation of unilamellar liposomes [22-24]. Sodium cholate was used in this study because it is non-toxic at low concentration and is easy to remove by dialysis [25].

#### Radiolabelling of nanoemulsions

Nanoemulsions were either single- or double-labeled with radioactive cholate and phospholipid. Single labeling refers to the use of [^3^H]-labeled cholic acid and double labeling refers to the use of both [^3^H]-labeled cholic acid and [^14^C]-phosphatidylcholine. Single labeling was performed by an overnight incubation of [^3^H]-labeled cholic acid (55.5 kBq) with nanoemulsion. The resulting radiolabelled nanoemulsion was called hot nanoemulsion. For double labeling, [^3^H]-labeled cholic acid (55.5 kBq) was used in the aqueous phase and L-α-dipalmitoyl-[^14^C]-phosphatidylcholine (DPPC) (5.55 kBq) was used in the oil phase. Thus obtained hot crude emulsion was extruded as usual. The initial concentrations of sodium cholate and phospholipid were 30 mM and 15.75 mM, respectively.

### Removal of sodium cholate by dialysis

After extrusion, the detergent was removed via dialysis through a very high permeability membrane (10 kDa cutoff). The nanoemulsion was dialyzed against a detergent free medium (glycerol, 2.5 % w/w) using an in-house built dialyzer (Fig 1) and a commercial dialyzer (Slide-A-Lyzer^®^).

#### Dialysis using in-house built dialyzer

The dialyzer was developed at an institute of Albert Ludwig University of Freiburg, Germany. The membrane (very high permeability 10 kDA cutoff membrane, Dianorm GmbH, Munich, Germany) was equilibrated with the dialysis fluid (glycerol, 2.5 % w/w) for at least 30 min prior to use and fixed in between the two compartments of a dialysis cell (4.9 cm^2^ cavity area) and the dialysis was performed as described elsewhere [26]. Briefly, a specified volume of nanoemulsion was placed in one compartment and the dialysis fluid was allowed to flow continuously through the other compartment. Both the compartments equipped with small magnetic stir bars were separated by the pre-equilibrated membrane. Any air bubbles in both the compartments were excluded to ensure enough osmotic pressure for detergent removal. Failure to seal the compartments tightly might result into loss of sample due to leakage. The flow rate (2.5 mL·min^-1^) of dialysis fluid was regulated by using a pump (Ismatec SA, Zurich, Switzerland). The dialysis was performed at continuous stirring (700 rpm) for 28 h at room temperature.

***Fig 1. In-house built dialyzer***

#### Dialysis using commercial dialyzer

Dialysis was performed by using a commercial dialyzer (Slide-A-Lyzer^®^) available in different capacities (0.5 mL – 3.0 mL) and membrane cutoffs. The dialyzer cassette (1 mL) having a membrane cutoff of 10 kDa was pre-equilibrated with the dialysis fluid (glycerol, 2.5 % w/w) for at least 30 min prior to dialysis. Nanoemulsion (1 mL) was pipetted into the cassette and a small magnetic stir bar was inserted into the cassette. Entrapped air was removed by lightly pressing the membrane and immediately closing the lid. Nanoemulsion was dialyzed against the fixed volume of dialysis fluid (500 ml) at constant stirring (300 rpm) for 28 h at room temperature.

#### Dialysis of radiolabelled nanoemulsions

Efficiency of dialysis was investigated for radiolabeled nanoemulsions dialyzed using two types of dialyzers. The radioactivity [^3^H and/or ^14^C] in nanoemulsions was analyzed by using liquid scintillation counter (LSC). As negative and positive controls, cold and hot nanoemulsions (before dialysis), respectively, were used. The negative control represented the background value, whereas, the positive control represented the reference value. For LSC measurement, the samples were withdrawn hourly, diluted with Ultima Gold^®^ at a ratio of 1:6 and analyzed under LSC to detect the radioactivity. All the measurements were performed in triplicates.

### Characterization of nanoemulsion

Nanoemulsions were characterized by measuring the particle size (Z-average), size distribution (polydispersity index, PDI) and the surface charge (zeta-potential) using a photon correlation spectroscopy (PCS, Malvern Nano ZS® series, Malvern, UK) which is based on Mie scattering theory [7]. A monochromatic laser 633 nm, fixed at a scattering angle of 173° is used to measure the Brownian motion of the particles which is correlated with the hydrodynamic diameter. Morphological characterization of nanoemulsions was performed by the help of cryo-transmission electron microscopic (cryo-TEM) pictures. The phospholipid content of the nanoemulsions was quantified by Bartlett assay [27]. Similarly, other parameters such as stability were also investigated.

#### Z-average and polydispersity index (PDI)

The z-average (nm) and polydispersity index (PDI) which represent the hydrodynamic diameter and size distribution of a particle, respectively was calculated as an average value of 3 consecutive measurements each consisting of 15 sub-runs lasting 10 s per sub-run. For the measurement, samples were prepared in a small volume disposable cuvette by diluting 5 µL nanoemulsion with 995 µL particle free deionized water (1:200). Prior to dilution deionized water was filtered through a cellulose acetate filter (Minisart^®^, Sartorius Stedim Biotech GmbH, Goettingen, Germany, pore size 0.2 µm) to avoid the effects of multiple scattering from dust particles. During the measurement, an equilibration period of 80 s and temperature of 25 °C were set up. The intensity average diameter and PDI of each sample was calculated from the Cumulant analysis (Zetasizer software 6.2) of each sample’s correlation function. The PDI indicates the homogeneity of the particle size distribution. A PDI value below 0.1 is an indication for a narrow size distribution [28].

#### Zeta potential (ζ)

The zeta-potential (ζ) is a charge acquired by a particle or molecule in a given medium and is measured by laser Doppler anemometry (Malvern Nano ZS^®^ series, Malvern, UK). Samples were diluted (1:200) similar to that for the particle size measurement and filled in a folded capillary cell. After an equilibration period of 120 s at 25 °C, measurements were performed in triplicate in an automatic mode so that the total sub-runs are between 10 and 100. Since the similar charges repel each other, the particles avoid phenomena such as flocculation and aggregation making the samples stable for longer period. Therefore, measurement of zeta-potential is an important parameter to study the stability of colloidal systems. Absolute values larger than ± 30 mV are considered as an indicator for a stable emulsion system [7].

### Cryo-transmission electron microscopy (Cryo-TEM)

Cryo-TEM is a widely used method to morphologically characterize the colloidal particles such as liposomes [26] and nanoemulsions [29]. This advanced microscopic method captures the two dimensional image of the sample and gives accurate information about the size, lamellarity and size distribution. The images of nanoemulsions were taken using a LEO 912 OMEGA electron microscope (Zeiss, Oberkochen, Germany) operating at 120 kV and ‘zero-loss’ conditions. Approximately 5 µL of sample (diluted if necessary) was placed on a copper grid (Quantifoil® S7/2 Cu 400 mesh, holey carbon films, Quantifoil Micro Tools GmbH, Jena, Germany) and any excess liquid was absorbed by a filter paper, so that only a thin (100 – 500 nm) liquid film remained on the copper grid [30]. The sample was then immediately shock-frozen by plunging it into liquid ethane. The vitrified sample was stored at 90 K (−183° C) in liquid nitrogen until it was loaded into a cryogenic sample holder (D626, Gatan Inc, Pleasanton, USA). The specimens were examined at −174 °C. Digital images were recorded with a slow scan CCD camera system (Proscan HSC 2 Oxford instruments, Abingdon, USA), and at a minimal under-focus of the microscope objective lens to provide sufficient phase contrast [31]. All the pictures were analysed using the software iTEM 5.0 Build 1054 (Soft Imaging System GmbH, Muenster, Germany). Various scales (2 µm, 1 µm, 500 nm, 200 nm, 100 nm) could be used to estimate the size of individual particle.

### Determination of phospholipid content

Phospholipid used in the preparation was quantified by performing phosphorous assay. This assay measures the phosphorous present in the head region of phospholipid as a phosphate molecule. The assay was performed according to the previously established method with some modifications [27]. The principle behind this assay is that one phosphorous atom corresponds to one phospholipid molecule.

The complete assay was conducted in phosphate free glass tubes. For a calibration curve, a standard solution of KH_2_PO_4_ (1 mM) in HCl (0.05 N) was prepared and the volumes of 50 µL, 100 µL, 150 µL, 200 µL, 250 µL, 300 µL and 350 µL were weighed in glass tubes on the assumption of Lambert-Beer law that absorbance is linear with concentration. Similarly, the sample volume was calculated from the theoretical concentration so that the phosphate content falls within the calibration curve. An empty glass tube (without standard solution) was used as blank value and treated in a similar manner as calibration and sample tubes. Then 500 µL of H_2_SO_4_ (10 N) were added to all the tubes including the blank tube and mixed well by vortexing and incubated at 160 °C for 3 h. After 3 h, the samples appeared dark brown in color due to oxidation. In order to completely oxidize the organic compounds, 200 µL of H_2_O_2_ (30 % w/w) was added, vortexed and incubated further at 160 °C for 1.5 h. Upon complete incubation, the solution in the tubes should turn clear. If this was not the case, incubation at 160 °C for 1.5 h was repeated with additional 200 µL of H_2_O_2_ (30 % w/w). Clear solutions marked complete oxidation and were ready for reduction process.

Then 4.75 mL of ammonium molybdate solution (0.22 % w/v) and 200 µL of freshly prepared Fiske Subbarrow reducer solution (14.8 % w/v) were added. After each addition of reagents, the contents of the glass tubes were mixed properly by vortexing. Then the tubes were covered with glass marbles and incubated for another 10 min at 95 °C in a heating block (MTB 250, Development and Technology, Ilmenau, Germany). After the incubation, the tubes were cooled down and vortexed. The solutions were transferred to a 2 mL disposable plastic cuvette and absorbance was measured using a spectrophotometer at a wavelength of 833 nm (Lambda XLS, Perkin Elmer, Hamburg, Germany). The blank value was deducted from the standard solutions and a calibration curve was prepared by plotting absorbance against the amount of phospholipid (micromoles). Using the slope of a straight line, the phospholipid concentration of the sample was calculated. The acceptable regression value of calibration curve was greater than 0.99.

### Stability studies

Nanoemulsions were stored under refrigeration for as long as 7 months. The particle size, PDI and zeta-potential were measured every month and compared with the initial value (day of preparation). Any significant increase in the above mentioned parameters was considered to be an unstable preparation. In addition, any phase separation if observed during the storage period was noted and concluded to fail the stability study.

## Results and discussion

### Characterization of nanoemulsion

#### Hand extruded nanoemulsions

The z-average of nanoemulsions after four sets of extrusion (51×400 nm and 153×200 nm membranes) was measured and the results are presented in Table 2. The z-average was found to be about 235 nm with PDI of about 0.135 after the first set of extrusion. Further extrusion through 200 nm membrane resulted in a sharp reduction in droplet size to about 185 nm with PDI below 0.09. Similarly, after the four sets of extrusion, the particle size reduced to about 166 nm and the PDI value was much lower (about 0.05). The size of emulsion droplet is smaller than the pore size of membrane (200 nm) because the droplets break down into droplet size closer to the pore size of the membrane [32]. Thus, it was observed that the reduction in droplet size can be improved by increasing the number of extrusion cycles. In case of liposomes, extrusion could reduce the vesicles to be in the size range between 50 and 100 nm when extruded through 100 nm pore size membranes [33]. However, the use of 100 nm membrane was not able to reduce the emulsion droplet below 100 nm probably due to the oily inner core (data not shown). The PDI value between 0.04 and 0.08 is defined to be extremely highly mono-disperse [28]. Therefore, the resulting nanoemulsion could also be considered as extremely mono-disperse nanoemulsions.

**Table 2.**
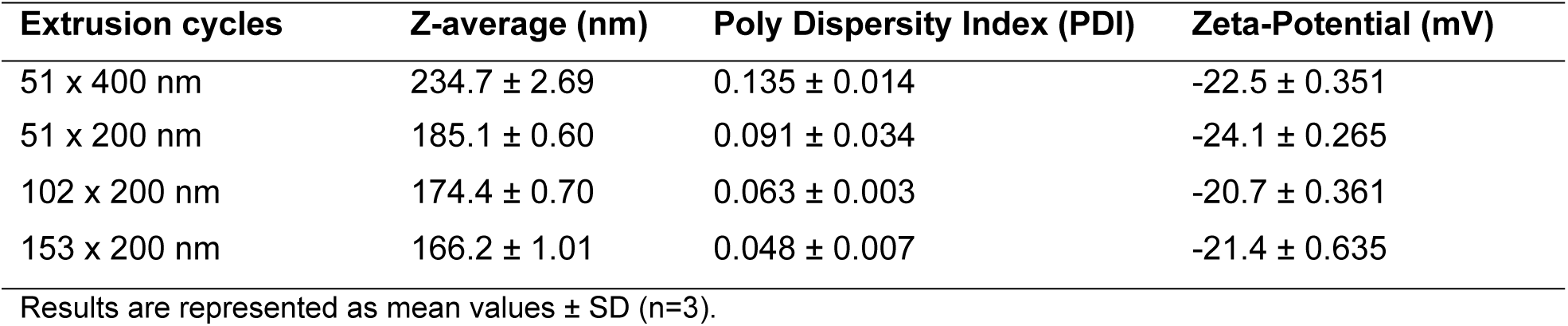
Characteristics of nanoemulsions after subsequent sets of extrusion cycles.

| Extrusion cycles | Z-average (nm) | Poly Dispersity Index (PDI) | Zeta-Potential (mV) |
| --- | --- | --- | --- |
| 51 x 400 nm | 234.7 ± 2.69 | 0.135 ± 0.014 | -22.5 ± 0.351 |
| 51 x 200 nm | 185.1 ± 0.60 | 0.091 ± 0.034 | -24.1 ± 0.265 |
| 102 x 200 nm | 174.4 ± 0.70 | 0.063 ± 0.003 | -20.7 ± 0.361 |
| 153 x 200 nm | 166.2 ± 1.01 | 0.048 ± 0.007 | -21.4 ± 0.635 |

Results are represented as mean values ± SD (n=3).

The zeta-potential of the emulsions during the various extrusion steps was found to be between −20 and −25 mV (Table 2). According to Hippalgaonkar and group, the obtained nanoemulsions are not electrostatically stable as the zeta-potential is below ± 30 mV [7]. It was also observed that the zeta-potential was not affected by increasing the extrusion cycle.

#### Nanoemulsions with sodium cholate

Since the sodium cholate was used only to disorganize the liposomal membrane [19], it was removed after extrusion by dialysis against detergent-free aqueous medium [25, 26]. Sodium cholate at various concentrations (20 – 50 mM) was studied for optimizing the appropriate concentration of sodium cholate to increase the number of nanoemulsions and reduce the number of liposomes. It was observed that the particle size changed minimally with an increase in sodium cholate concentration (Table 3).

**Table 3.** Characteristics of nanoemulsions with different concentrations of sodium cholate.

| Sodium Cholate (mM) | Z-average (nm) |  | Poly Dispersity Index (PDI) |  | Zeta-Potential (mV) |  |
| --- | --- | --- | --- | --- | --- | --- |
|  | Before Dialysis | After Dialysis | Before Dialysis | After Dialysis | Before Dialysis | After Dialysis |
| 20 | 161.6 ± 1.44 | 160.8 ± 2.69 | 0.079 ± 0.025 | 0.072 ± 0.018 | -45.4 ± 6.11 | -35.8 ± 16.7 |
| 30 | 168.6 ± 10.32 | 165.9 ± 8.90 | 0.050 ± 0.006 | 0.056 ± 0.004 | -47.3 ± 2.57 | -37.8 ± 6.7 |
| 40 | 165.9 ± 11.66 | 166.1 ± 9.12 | 0.023 ± 0.017 | 0.069 ± 0.005 | -47.9 ± 7.95 | -36.1 ± 4.6 |
| 50 | 168.9 ± 12.05 | NA | 0.043 ± 0.001 | NA | -44.4 ± 3.03 | NA |

Extrusion cycles: 51x400 nm and 153x200 nm, Results are represented as mean values ± SD (n=3). NA: Not available.

However, the PDI value was found to remain below 0.1 at all concentrations of sodium cholate. After complete extrusion (51×400 nm and 153×200 nm), the size of nanoemulsions was found to be below 170 nm at all concentrations of sodium cholate which is comparable to the size of nanoemulsions without sodium cholate (Table 2). Additionally, dialysis seemed to have negligible effect on particle size and PDI of nanoemulsions. Thus, the findings suggest that all the studied concentration of sodium cholate (20 – 50 mM) can be used to prepare homogenous nanoemulsions without affecting the Z-average and PDI values.

Similarly, an increase in sodium cholate concentration did not increase the zeta-potential of nanoemulsions (Table 3).The zeta-potential was found to be in the range of −45 mV to −48 mV when sodium cholate was used at different concentrations (Table 3) which is an increment of about −20 mV when compared to that of nanoemulsions without sodium cholate (Table 2). Nevertheless, after dialysis the zeta-potential was reduced to about −36 mV which accounts to a loss of about −10 mV. Since the values were above −30 mV, the nanoemulsions were stable electrostatically. It was observed that the use of sodium cholate provided additional electrostatic stability to the nanoemulsion even after its removal via dialysis. Therefore, the use of sodium cholate at higher concentration (50 mM) does not seem to be beneficial.

### Removal of sodium cholate by dialysis

Two types of dialyzers were used to compare their detergent removal performance and the results are shown in Table 4.

**Table 4.** Comparison of dialyzers.

| Dialyzers | Z-average (nm) |  | Poly Dispersity Index (PDI) |  | Zeta-Potential (mV) |  |
| --- | --- | --- | --- | --- | --- | --- |
|  | Before Dialysis | After Dialysis | Before Dialysis | After Dialysis | Before Dialysis | After Dialysis |
| In-house dialyzer | 164.5 ± 7.31 | 165.1 ± 9.11 | 0.039 ± 0.006 | 0.058 ± 0.022 | -50.4 ± 4.15 | -43.5 ± 1.48 |
| Slide-A-Lyzer® | 165.5 ± 7.59 | 164.3 ± 7.96 | 0.049 ± 0.013 | 0.062 ± 0.006 | -49.47 ± 4.80 | -40.37 ± 2.39 |

Extrusion cycles: 51x400 nm and 153x200 nm, Sodium cholate concentration: 30 mM. Results are represented as mean values ± SD (n=3).

The concentration of sodium cholate was maintained at 30 mM. It was observed that the nanoemulsions before and after the dialysis were similar in size and homogeneity in both dialyzers. But the zeta-potential after dialysis using the Slide-A-Lyzer^®^ was found to be slightly lower than the in-house built dialyzer. However, the nanoemulsions were stable with zeta-potential above −30 mV in both the cases. These findings showed that the two dialyzers showed similar performance and are easily replaceable as per the convenience and availability.

#### Sodium cholate removal efficiency

Sodium cholate removal efficiency was investigated using the two types of dialyzers and the results are shown in Fig 2.

***Fig 2. Sodium cholate removal efficiency in radiolabelled nanoemulsions (single labeling)***

Labeling: Incubation of cold nanoemulsion with [^3^H]-labelled cholic acid (55.5 kBq). Sodium cholate: 30 mM. Results are normalized with a dilution factor calculated after dialysis. Error bars represent SD (n=3).

The study illustrated that sodium cholate removal profile differed slightly in the early phase but overlapped with each other in the later phase, showing similar pattern of detergent removal in both the dialyzers. Within the first hour of dialysis only about 16 % of cholate was removed by the Slide-A-Lyzer^®^ whereas already 58 % was removed by the in-house dialyzer. However, after 7 hours of dialysis, both the dialyzers exhibited similar efficiency (detergent removal of about 85 %). After the completion of dialysis period of 28 h, the residual cholate for both dialyzers was found to be between 3 and 5 % when the molar ratio of phospholipid-to-detergent was 0.525 (15.75 mM phospholipid and 30 mM sodium cholate). In a previous study, the residual cholate was measured to be less than 0.5% after 24 h of dialysis when the molar ratio of phosphatidylcholine-to-cholate was maintained at 0.625 [28]. This explains that the amount of residual detergent depends upon the phospholipid-to-detergent ratio used.

Similarly, in double labeling [^3^H and ^14^C] study, along with the sodium cholate depletion, phospholipid content was also analyzed to monitor the loss of phospholipid during dialysis. Radiolabelled nanoemulsion was dialyzed using Slide-A-Lyzer^®^. The samples were analyzed hourly for ^3^H and ^14^C under LSC and the results are summarised in Fig 3. It was observed that about 65 % of sodium cholate was removed within 2 h, whereas, after 28 h of dialysis, about 7 % of sodium cholate remained in nanoemulsion (Fig 3A). This value was slightly higher than the amount obtained in the previous study where labeling was performed by incubating cold nanoemulsion with [^3^H]-labelled cholic acid (Fig 2, Slide-A-Lyzer^®^). This study thus showed that the method of radiolabeling affects the sodium cholate removal profile. The ratio ^3^H/^14^C was about one which means that almost an equal proportion of ^3^H and ^14^C remained in the nanoemulsion after dialysis (Fig 3B).

***Fig 3. Analysis profile of nanoemulsion dialyed using Slide-A-Lyzer*^*®*^*(double labeled) A)* ^*3*^*H and* ^*14*^*C (Radioactivity %); B) Ratio of* ^*3*^*H and* ^*14*^*C (*^*3*^*H/*^*14*^*C)***

Labeling: Extrusion of hot crude emulsion containing [^3^H]-labelled cholic acid (55.5 kBq) and [^14^C]-Phosphatidylcholine (5.55 kBq). Sodium cholate: 30 mM. Results are normalized with a dilution factor calculated after dialysis. Results are represented as mean values. Error bars represent SD (n=3).

### Cryo-transmission electron microscopy (Cryo-TEM)

The cryo-TEM pictures were not only essential to observe the presence of liposomes but also to find out the appropriate concentration of sodium cholate required to prepare homogenous and stable nanoemulsions. The Cryo-TEM pictures of nanoemulsions at different conditions are shown in Fig 4.

***Fig 4. Cryo-TEM pictures of nanoemulsions at different conditions.***

***A) Without sodium cholate; B) 20 mM sodium cholate after dialysis; C) 30 mM sodium cholate after dialysis; D) 40 mM sodium cholate after dialysis; E) 30 mM sodium cholate before dialysis; F) 30 mM sodium cholate after dialysis and after 23 weeks of storage under refrigeration.***

Extrusion cycles: 51×400 nm and 153×200 nm. Dialyzer: in-house dialyzer.

It was surprisingly observed that very few nanoemulsion droplets (dark circular structures) and many liposomes (transparent circular structures) were present when sodium cholate was not used in the preparation of nanoemulsions (Fig 4A). In cryo-TEM images, liposomes appeared as transparent circular structures due to their aqueous interior and the dark border represents the phospholipid bilayer [26], whereas, nanoemulsions appeared as dark circular structures due to their oily inner core. According to Torchilin and Weissig, liposomes are formed spontaneously upon rehydration of phospholipids [34, 35]. Therefore, the phospholipid used as emulsifying agent in the preparation of nanoemulsions could also form liposomes. Previously, nanoemulsions and solid lipid nanoparticles were prepared by using extrusion method but without the use of sodium cholate. Also, the simultaneous formation of liposomes was not mentioned earlier [36]. Our findings suggest that cryo-TEM pictures are necessary in complete characterization of nanoemulsions. If cryo-TEM pictures are not taken, the presence of liposomes along with nanoemulsions could not be identified. From Table 3, it was difficult to find out the optimum concentration of sodium cholate to prepare homogenous and stable nanoemulsions which are free from liposomes because at all concentrations of sodium cholate, nanoemulsions were found to be homogeneous and stable on the basis of PDI and zeta-potential values. Therefore, cryo-TEM pictures of nanoemulsions were supportive to find out the optimum concentration of sodium cholate.

Fig 4 revealed that after dialysis, liposomes of about 200 nm (marked with white arrows) were present at sodium cholate concentration of 20 mM (Fig 4B), but at 30 mM, no such liposomes were detected (Fig 4C). As the concentration of sodium cholate was increased further to 40 mM, numerous but very small liposomes (marked with white boundaries) were observed again (Fig 4D). At 20 mM, the sodium cholate was perhaps adequate to solubilize the lipid membrane and reorganise them to form liposomes upon dialysis. But at 30 and 40 mM, the ratio of detergent-to-phospholipid was perhaps inadequate to form liposomes having defined size. In a previous study, the critical molar ratio of detergent-to-lipid for the formation of liposomes by detergent removal via dialysis was found to be between 1.2 and 2 with lipid up to 25 mM [20, 28]. With the help of cryo-TEM pictures, it was thus concluded that the appropriate concentration of sodium cholate to prepare nanoemulsions without liposomes was 30 mM.

The cryo-TEM picture of nanoemulsion before (Fig 4E) and after dialysis (Fig 4C) seem to be similar indicating no influence of dialysis on the size of nanoemulsion and the results were also supported by Table 3 and Table 4. Additionally, Fig 4C and Fig 4F do not seem to differ much which means that the nanoemulsion was stable for as long as 23 weeks when stored under refrigeration.

### Determination of phospholipid content

Phospholipid content of nanoemulsions prepared using 30 mM sodium cholate was quantified before and after dialysis by means of phosphorous assay. With 1.2 % (w/w) of phospholipid (E80^®^), the initial concentration was theoretically calculated to be 15.75 mM. After phosphorous assay, the phospholipid content was found to differ slightly (Table 5).

**Table 5.** Phosphorous assay of nanoemulsions.

| Type of dialyzer | Phospholipid content (mM) |  |
| --- | --- | --- |
|  | Before dialysis | After dialysis |
| In-house built dialyzer | 14.9 ± 2.38 | 11.73 ± 1.38 |
| Slide-A-Lyzer® | 14.54 ± 0.92 | 14.11 ± 1.62 |

It was observed that phospholipid was lost during dialysis. Among the two types of dialyzers, the loss was found to be higher in in-house dialyzer (21 %) than in Slide-A-Lyzer^®^ (3 %). The variation in loss of phospholipid in both dialyzers could be explained by the variation in dialysis conditions such as volume and flow rate of dialysis fluid. The dialysis process in in-house dialyzer was an open-system where the dialysis fluid (about 4.5 L) was allowed to flow continuously at a fixed flow rate (2.5 mL min^-1^) for a fixed period of time (28 h) whereas, dialysis using Slide-A-Lyzer^®^ was a closed-system where a fixed volume of dialysis fluid (500 mL) at constant stirring (300 rpm) was used and the dialysis for continued for 28 h at room temperature. From this, it is clear that the volume of dialysis fluid affects the loss of phospholipid but does not affect the characteristics of nanoemulsions.

In spite of the loss of phospholipid during dialysis (Table 5), the cryo-TEM images revealed that the nanoemulsions were still stable even after storage under refrigeration for as long as 23 weeks (Fig 4 F) when compared to the cryo-TEM images before dialysis (Fig 4 E) and after dialysis (Fig 4 C).

### Stability studies

The stability of nanoemulsions dialyzed by using two different dialyzers was studied for a duration of 7 months. Since the measurement of particle size, PDI and zeta-potential and Cryo-TEM images are useful techniques to confirm the stability of nanoemulsions [37], the samples were monitored every month for particle size, PDI and zeta-potential. Three samples per dialysis method were stored under refrigeration (4 – 8 °C) and studied for their stability. Fig 5 represents the summary of stability of nanoemulsions dialyzed using in-house built dialyzer.

***Fig 5. Stability of nanoemulsions dialyzed using in-house built dialyzer***

It was observed, that the changes in size during the storage period was negligible. The PDI values were found to change with time but remained below 0.1 even after 7 months of storage. A slight fluctuation was noted in zeta-potential for all samples. Nevertheless, zeta-potential was measured to be above −30 mV for all samples even after 7 months. Therefore, the samples were concluded to be stable under refrigeration for as long as 7 months.

Similarly, the stability of nanoemulsions dialyzed using Slide-A-Lyzer^®^ is summarized in Fig 6. As shown in figure, the changes in both size and PDI were observed to be negligible like in the case of in-house built dialyzer (Fig 5). The zeta-potential throughout the storage period was measured to be above −30 mV. Due to all these reasons, the samples were found to be stable for about 7 months. Therefore, both dialyzers were found to be suitable for preparing nanoemulsions without liposomes which are stable for as long as 7 months under refrigeration. Thus, both the dialyzers are conveniently replaceable to each other.

***Fig 6. Stability of nanoemulsions dialyzed using Slide-A-Lyzer*^*®*^**

## Conclusions

Preparation of homogenous nanoemulsions in small-scale is a challenge, especially if the emulsion is stabilized by phospholipids due to the unavoidable formation of liposomes along with the emulsion droplets. Extensive extrusion of crude emulsion through polycarbonate membranes (51× 400 nm and 153× 200 nm) at 65 °C not only reduced the emulsion droplet to nanometer range and but also prepared homogenous droplets as depicted by a PDI value below 0.1. Furthermore, the study showed that the use of a physiological detergent, sodium cholate at 30 mM concentration and later removal via dialysis after extrusion minimized the formation of liposomes resulting into nanoemulsions which are stable under refrigeration for as long as seven months. The cryo-TEM pictures provided sufficient evidences that the use of sodium cholate was indeed beneficial to prepare liposome-free nanoemulsions in small scale (less than 1 mL). The easy availability of commercial dialyzers at variable capacities makes the preparation process even easier.

Thus, this method could be regarded as an economic and yet promising technique especially for preparing functionalized or modified nanoemulsions where expensive ligands or antibodies, fluorescence or radioactive markers must be used to target such nanoemulsions to a specific cell or location.

## Acknowledgements

The authors are thankful to DAAD for providing scholarship to conduct this project and Lipoid for the generous gift of phospholipid.

## Author contributions

Conceived and designed the experiments: SG and RS. Performed the experiments: SG and SB. Analyzed the data: SG, MZ and RS. Contributed reagents/materials/analysis tools: SG, MZ and SB. Wrote the paper: SG, MZ, SB and RS.

